# Transcription-dependent swelling of a transplanted chromosome in an artificial cell

**DOI:** 10.1101/2024.09.25.614905

**Authors:** Ferdinand Greiss, Shirley S. Daube, Vincent Noireaux, Roy Bar-Ziv

**Affiliations:** Department of Chemical and Biological Physics, Weizmann Institute of Science, Rehovot, 7610001, Israel; School of Physics and Astronomy, University of Minnesota, Minneapolis, MN, 55455, USA

**Keywords:** Synthetic biology, Biophysics, Cell-free chromosomes, Cell-free gene expression, Genome organization

## Abstract

Transplanting chromosomes from living to artificial cells would impact our understanding of chromosome organization and DNA transactions, with implications for autonomous biological systems. Here, we transplanted *Escherichia coli* chromosomes into artificial cells, enabling real-time labeling, manipulation, and steady-state gene expression down to the single-molecule limit. Chromosomes stripped of native proteins transitioned from a swollen to compacted state induced by transcription inhibition, in contrast to protein-bound chromosomes retaining an organization with blobs. In a cell-free expression reaction, RNA polymerases were uniformly distributed along the entire chromosome and rapidly detached, consistent with a global transcriptional activity. We used tailored surfaces to capture and count 20 nascent proteins per hour from a single gene on the chromosome. We mapped stably bound condensins to the blobs, supporting a model where swelling by transcription is counterbalanced by condensin-mediated compaction. Our data suggest transplanted chromosomes as active gels organized by molecular machines.

## Main text

Programming artificial cells with single chromosomes presents the next frontier for bottom-up synthetic biology. Chromosomes have been transplanted from one species to another (1) and chemically synthesized and assembled from smaller fragments (2–5), functioning as the genetic blueprint for living cells. The bacterial genome, such as *E. coli*, is encoded on a more than 10^6^ base-pair long contiguous polymer, packed from its contour length of a few millimeters into the 1000-fold smaller cell. The long DNA molecules dramatically expand when released from cells (6,7). Active and passive processes condense the long DNA molecule into 3-D organizations (8–10), compact yet amenable to dynamic opening processes (11–13), but the direct link between DNA organization and DNA transactions remains an outstanding question. Transplanting into artificial cells could advance our understanding by releasing the chromosome from its confined small volume and complex intertwined dynamics *in vivo*. Further engineering – with the required knowledge of such fundamental processes - could establish chromosomes as active material for future synthetic cells (14).

Single-molecule studies *in vitro* have advanced our understanding of isolated DNA transactions with relatively short DNA molecules (15–22), whereas engineering an active system supporting a broad spectrum of biochemical processes requires reconstituting the entire flow of genetic information at the genomic scale. Although powerful, current methods on cell-free chromosomes using closed settings (23–30), such as encapsulated cell mimics, e.g., liposomes, have limitations on exchanging solutions, external manipulation, and imaging. The two-step process of isolating and later embedding a chromosome also increases the risk of breakage of such long DNA molecules. The challenge lies in integrating individual chromosomes with *in vitro* steady-state conditions for cell-free gene expression, allowing real-time observation of proteins with low abundance and manipulation of numerous biological machines active on the DNA concurrently.

Here, we developed flat, semi-open compartments integrated into a multifunctional microfluidic chip as artificial cells to achieve holistic steady-state gene expression dynamics with single-molecule resolution. *E. coli* bacteria were trapped in compartments to extract their 4.6-Mbps long chromosomes through gentle *in situ* cell lysis, embedding a natural genome inside artificial settings either with (proteinated state) or without (deproteinated state) chromosome-bound proteins. Using high-resolution fluorescence microscopy, we followed cell-free gene expression upon the addition of an *E. coli*-based transcription-translation system and studying RNA polymerases (RNAP), ribosomes, and the bacterial condensin MukBEF to find transcription as an energy source that globally swells chromosomes, counterbalanced by local compacting MukBEF clusters.

### Transcription swells chromosomes in cell lysate

We constructed large diamond-shaped compartments (W=100 μm, H=70 μm) with a height of 1.5 μm. Two 75 μm deep flow channels flanked the compartment for continuous fluid exchange (**Fig. 1a and Supplementary Fig. 1a, b**). We loaded on the chip *E. coli* K-12 MG1655 bacteria, which were grown to the exponential phase and incubated with a sucrose buffer as a pre-conditioning step before cell lysis (6). After on-chip cell wall degradation, we exchanged buffers to lyse the cells *in situ*, gently transplanting the chromosome from the bacteria into the compartment, followed by incubation with the non-specific protease Proteinase K to remove chromosome-bound proteins (Methods). With the influx of a lysate-based *E. coli* cell-free transcription-translation system (prepared from the strain BL21) and supplemented with the intercalating DNA dye SYBR Green I (SG-I), we could observe chromosomes self-organizing into an intricate arrangement, likely due to steric hindrance by the neighboring chromosomes (**Fig. 1b, Supplementary Fig. 1c and d**). Most strikingly, we found that exchange to a cell lysate supplemented with 500 nM rifampicin, an inhibitor of transcription initiation, led to the compaction of all chromosomes within minutes (**Fig. 1b-d**), most likely induced by macromolecular crowding (6). The compaction was marked by a ∼2.7-fold average increase in SG-I signals and an estimated ∼50% area drop (**Fig. 1b-e, Supplementary Fig. 1c-f**). Because of the fuzzy boundaries of the swollen chromosomes, we found the area estimation error-prone and continued with the increased SG-I signal as a robust measure for chromosome compaction. We corroborated the compaction phenomenon by reproducing the experiments with a cell-free gene expression system prepared from another *E. coli* strain, the standard laboratory strain K-12 MG1655 (**Supplementary Fig. 1c-f**).

**Figure 1:**
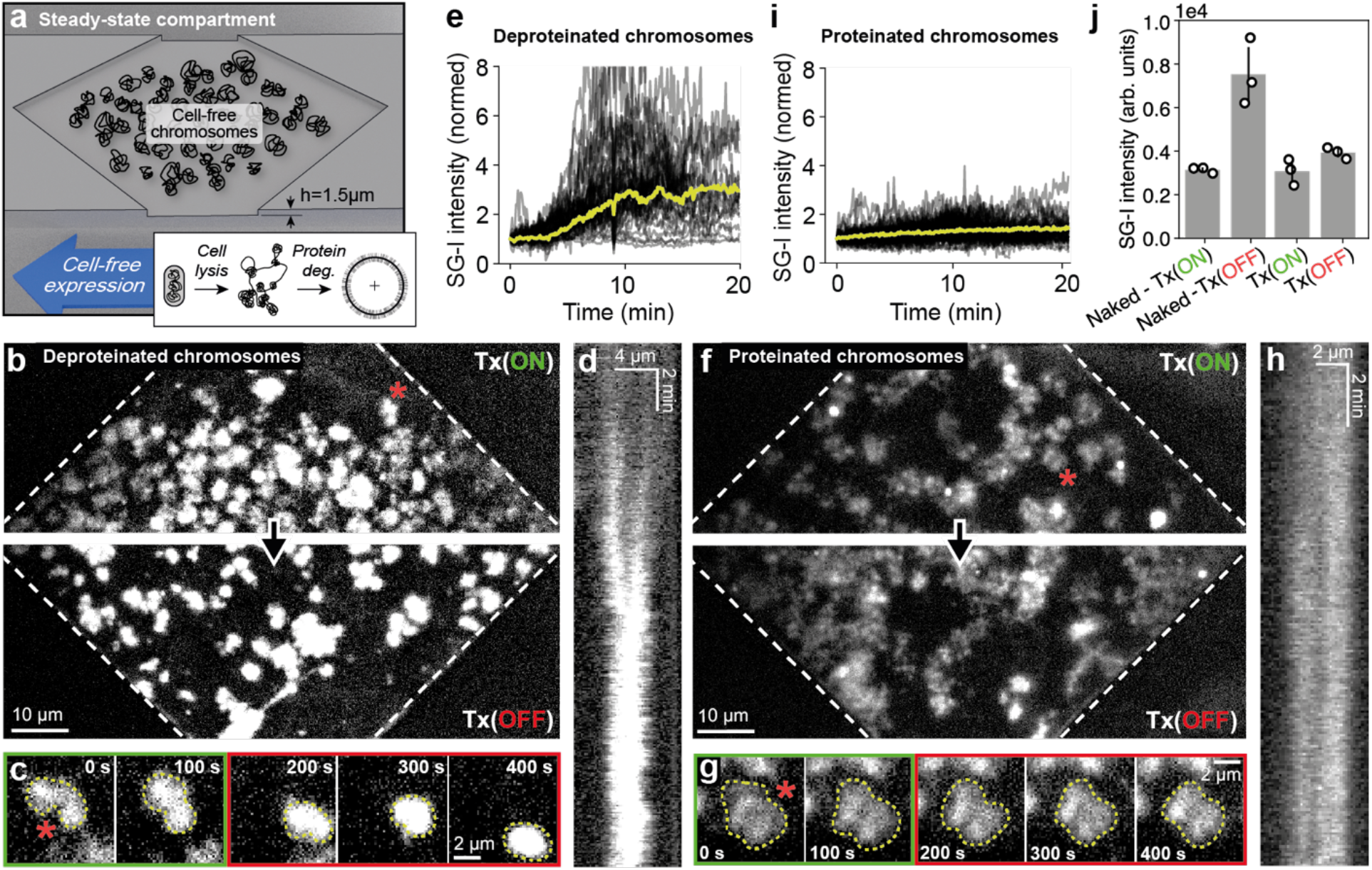
Transcription swells chromosomes *in vitro*. **a)** A 3-D rendering of a large diamond-shaped compartment flanked by two main flow channels to introduce *E. coli* cells, lysis buffer, and the cell-free gene expression system. **b)** A representative fluorescence image of deproteinated chromosomes transplanted from *E. coli* cells into the diamond-shaped compartment with a cell-free expression system prepared from *E. coli* BL21 Rosetta 2 under active (top image) and inhibited (bottom image) transcription (Tx), controlled by adding 500 nM of the transcription inhibitor rifampicin (Rif). Deproteinated chromosomes were produced through protein degradation with Proteinase K during the cell lysis step. The chromosomes were fluorescently labeled with the SYBR Green I (SG-I) intercalating DNA dye. **c)** A fluorescence time-lapse montage with a zoom on a deproteinated chromosome transitioning from active (green box) to inhibited (red box) Tx (the red asterisk annotates the location of the chromosome in panel b). The Tx activity in cell lysate was inhibited at ∼200s by the inflow of cell lysate incubated with 500 nM Rif. **d)** A kymograph of an SG-I labeled chromosome during the transition from active to inhibited Tx. **e)** The average (yellow curve) and individual (black curves) SG-I intensities of deproteinated chromosomes transitioning from active (up to ∼3 minutes) to inhibited (from 3 minutes onwards) Tx. **f)** Same as panel b for proteinated chromosomes. **g)** Same as in panel c for proteinated chromosomes. **h)** Same as in panel d for proteinated chromosomes. **i)** Same as in panel e for proteinated chromosomes. **j)** The average SG-I intensities before and after Tx turned off for deproteinated (naked) and proteinated chromosomes. The bars and error bars show averages and S.D. values of three independent experiments (black circles) done with two cell-free gene expression system preparations.

The observation that transcription inhibition led to the compaction of chromosomes was surprising, especially since the contrary was observed in live bacteria, where transcription inhibition by rifampicin has been shown to lead to transient compaction followed by chromosome expansion filling the ∼1 μm^3^ cell volume after tens of minutes (31–33). To investigate this discrepancy, we reasoned that chromosomes in their natural cellular environment are bound by many proteins, while our *in vitro* observation was made with deproteinated chromosomes. Therefore, we repeated the transcription-inhibition experiment without degrading the chromosome-bound proteins. Upon transcription inhibition, the proteinated chromosomes appeared less homogenous with blob-like structures (**Fig. 1f-h)** and displayed only a minor (∼1.3-fold) increase in the overall SG-I intensity (**Fig. 1i, j**). These results better reproduced the *in vivo* data, suggesting that chromosome-bound proteins prevented the global compaction.

To demonstrate that native proteins remained bound to the chromosomes during the transplantation process and that deproteination was effective, we performed a control experiment in which chromosomes were transferred from bacteria expressing *E. coli* RNA polymerase fused to the fluorescent HaloTag (HT) reporter system (fused to the gene *rpoC*, coding for the RNAP protein β’) (Methods, **Supplementary Fig. 2**) (34). Upon cell lysis, the chromosomes could be imaged by the RNAP fluorescent signal without adding SG-I, suggesting that the chromosomes retained at least some of their native proteins. Under these conditions, the chromosomes underwent an abrupt area increase directly after lysis, followed by a slower expansion (**Supplementary Fig. 2c, top row**), suggesting that a relaxed final chromosome state had been reached after a few minutes. With the addition of Proteinase K, the chromosomes lost the RNAP signal after cell lysis (**Supplementary Fig. 2b-d**), demonstrating that the deproteination of the chromosomes was efficient.

### Genome organization of a cell-free chromosome

We hypothesized that the spatial DNA organization, which seemed to have been maintained to some extent by the proteinated chromosomes, may have prevented the global compaction induced by transcription inhibition. To gain support, we sought to isolate individual proteinated chromosomes into smaller cell-like compartments (35–38) and investigate their DNA organization before cell-free gene expression was initiated. We constructed the compartments with 1.5 μm height and 20 μm diameter connected by two capillaries with the two flanking main flow channels (**Fig. 2 and Supplementary Fig. 3**). Bacteria (h∼1 μm) were centrifuged from the flow channel into the compartment through the thicker capillary (h∼1.5 μm) and were retained by the thinner constriction capillary (h∼400 nm). The embedded cells were lysed as previously. The cells contained a plasmid coding for the histone-like protein HUα fused to a green fluorescent protein (GFP), which binds DNA as nucleoid-associated protein with no strong sequence specificity (39,40) and is often used as a DNA labeling method (**Supplementary Fig. 4**).

**Figure 2:**
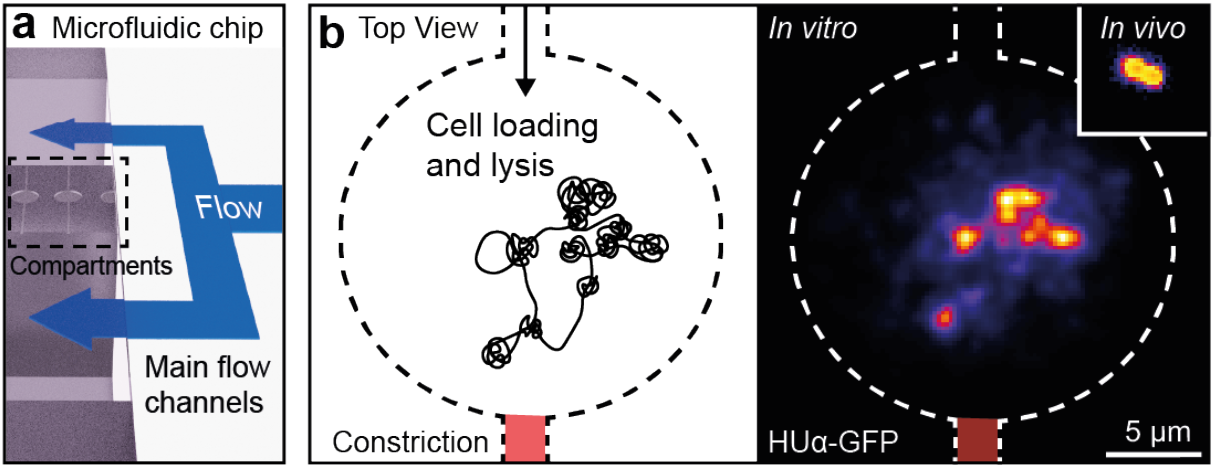
A chromosome embedded into a cell-like compartment. **a)** A 3-D rendering of the microfluidic chip with a cell-like compartment flanked by two main flow channels to introduce cells, lysis buffer, and a cell-free expression system. **b)** A schematic of a cell-like compartment. Cells were introduced from the top and retained by a thin capillary (constriction) during centrifugation of the microfluidic chip loaded with a cell suspension in sucrose buffer. The right panel shows a bacterium before cell lysis (inset) and a chromosome after cell lysis. The chromosome was labeled with HUα-GFP expressed from plasmids. The fluorescence image after lysis was deconvolved to highlight the blob-like structures. The scale bar applies to both images.

Upon *in situ* cell lysis, the transplanted HUα-GFP labeled chromosome appeared as a blob-like structure (**Fig. 2b**). We then applied a weak alternating electric field across the cell-like compartments to pull and release chromosome parts into and from the thin capillary (**Fig. 3a, b**). The HUα-labeled chromosome stretched and relaxed reversibly in about a second (**Fig. 3c**), supporting the notion that the blobs were stable structures along the intact chromosome (**Fig. 3a, Supplementary Fig. 5a**).

**Figure 3:**
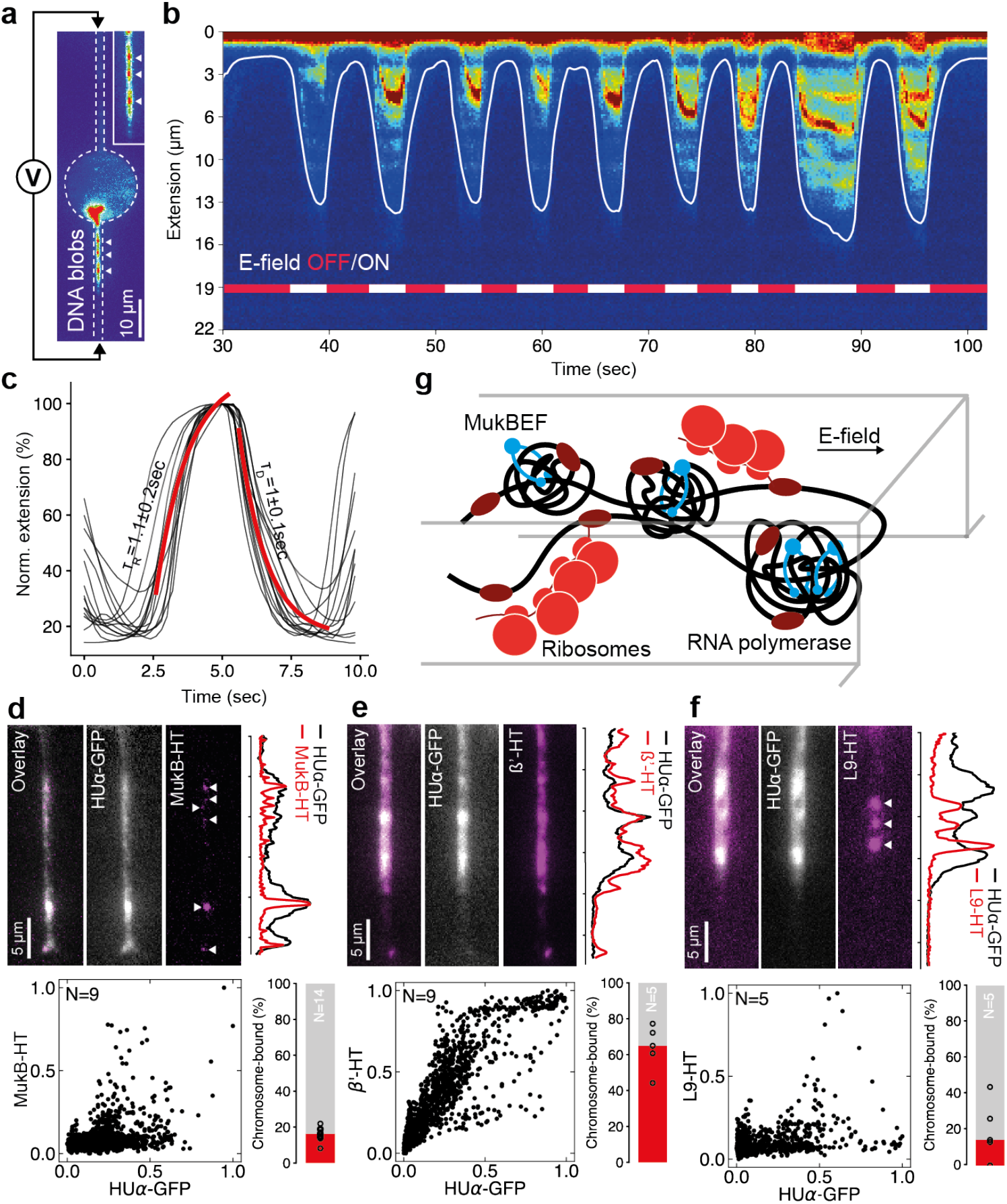
The spatial organization of molecular machines on a proteinated chromosome. **a)** Two electrodes were installed to apply an electric field across the cell-like compartment with a proteinated chromosome. Short chromosome segments were pulled into the capillary, revealing bright fluorescent HUα-GFP blobs (white arrowheads). **b)** An alternating electric field was applied along the compartment, and the HUα-GFP signal was measured as a line profile along the lower capillary. The continuous white line indicates the DNA boundary detected by image processing. **c)** The stretching and relaxing dynamics of the chromosome with the electric field turned on and off, respectively. The black curves indicate the repetitions and the red curves show global fits to mono-exponential raise and decay. **d)** Representative fluorescence snapshots of stretched segments from proteinated chromosomes. HUα-GFP was produced from plasmids, and *mukB* was chromosomally tagged with the HaloTag (HT). The double-color line profiles were measured on the capillary and normalized. The fluorescence signal showed a clustered localization of MukB-HT (white arrowheads), with the scatter data showing maximal values at intermediate (∼0.5) and maximal (1.0) HUα-GFP values. The scatter data was combined from several independent capillaries (the specific number is indicated in the graph). The fraction of chromosome-bound MukB-HT was computed from its signal before and after cell lysis where the red bar indicates the median from different cells (black circles). **e)** Same as panel d with a linear trend between the β’-HT and HUα-GFP signals. **f)** Same as panel d with high and low L9-HT signals at intermediate HUα-GFP signals and anti-correlated at higher HUα-GFP signals. **g)** A sketch to show the spatial organization of the three biological machines, MukBEF, RNAP, and ribosomes, along the stretched bacterial chromosome, as deduced from the results in panels d-f.

To investigate the content of these dense DNA blobs, we focused on the bacterial condensin complex MukBEF, proposed as an active loop extruder and key genome organizer that localizes across the chromosome as discrete clusters (41–45), each consisting of ∼36 MukB proteins (46) that entrap DNA locally. *In vivo* experiments reported a DNA-bound fraction of ∼20% with a short residence time of ∼50 seconds for the MukB on DNA (46). We tagged the *mukB* gene on the chromosome with HT (43), and despite losing the expected ∼85% of the MukB-HT signal during the *in situ* cell lysis due to unbound MukB (**Supplementary Fig. 4a, b**), we could co-localize the MukB and the HUα-GFP signals, demonstrating that the ∼15% MukB complexes remained bound and co-localized with the chromosome blobs (**Fig. 3d, Supplementary Fig. 4c, and Supplementary Fig. 5b**), and suggesting that MukB may be responsible for their formation.

Whether these blobs influence gene expression is an open question. We examined chromosomes from bacteria with the HT-labeled RNAP and HT-labeled ribosomes (fused to the gene *rpII*, coding for the ribosomal protein L9) (47) (**Supplementary Fig. 4a, b**). Using the RNAP-labeled *E. coli* cells, the chromosome entered the capillary with the RNAP signal covering it continuously, apparently unaffected by the different DNA densities (**Fig. 3e, Supplementary Fig. 5b**). In contrast, only a sub-fraction of the ribosome signal appeared as sparse bright foci (**Fig. 3f, Supplementary Fig. 5b**). About 65% of the RNAP and only ∼10% of the ribosome was retained on the chromosome after lysis (**Fig. 3e, f, respectively, and Supplementary Fig. 4c**). The ribosome foci were outside the blob-like DNA structures, as evident by the anti-correlation with the HUα-GFP signal, suggesting large polysome formation because of the strong fluorescent signal on DNA excluded from the local dense DNA environment. While the genome organization might, therefore, affect the small ribosome fraction involved in the formation of chromosome-bound polysomes, we concluded that gene expression, particularly transcription, seemed mostly independent of the genome organization (**Fig. 3g**).

### MukB activity during cell-free gene expression

Having identified MukB as a potential factor responsible for the *in vitro* chromosome organization, we repeated the cell-free gene expression experiments in the large compartments, but this time with fluorescently labeled MukB-HT from the source bacteria. Upon flushing in the cell-free gene expression system, we found MukB stably bound to the proteinated chromosomes during active transcription with very little turnover (**Fig. 4a, Supplementary Fig. 6a-c**). Despite minor HT labeling heterogeneity, the SG-I DNA signal accumulated on MukB-HT spots with a high colocalized fraction of ∼90% (**Supplementary Fig. 6d, e**), supporting the notion that MukB produced stable high-density DNA regions on the chromosome. To assess the relevance of MukB to chromosome swelling, we then measured changes in SG-I signal intensity upon transcription inhibition in the absence of only MukB using *E. coli* chromosomes with deleted *mukB* gene (43). In this case, the SG-I signal increased by an average of 1.7-fold with proteinated chromosomes, which is an intermediate value between the ∼1.3-fold and ∼2.7-fold compaction with proteinated and deproteinated K-12 MG1655 chromosomes, respectively (**compare Fig. 4b, c to Fig. 1 b, d, respectively**). The combined data suggested that MukB is a critical player in the effects imposed by transcriptional activities and macromolecular crowding. We propose a model with three major processes that act on a cell-free bacterial chromosome: Transcription as a source for genome-scale extensile swelling, MukBEF as a source for point-like compaction, and macromolecular crowding as a global compaction force (**Fig. 4d**).

**Figure 4:**
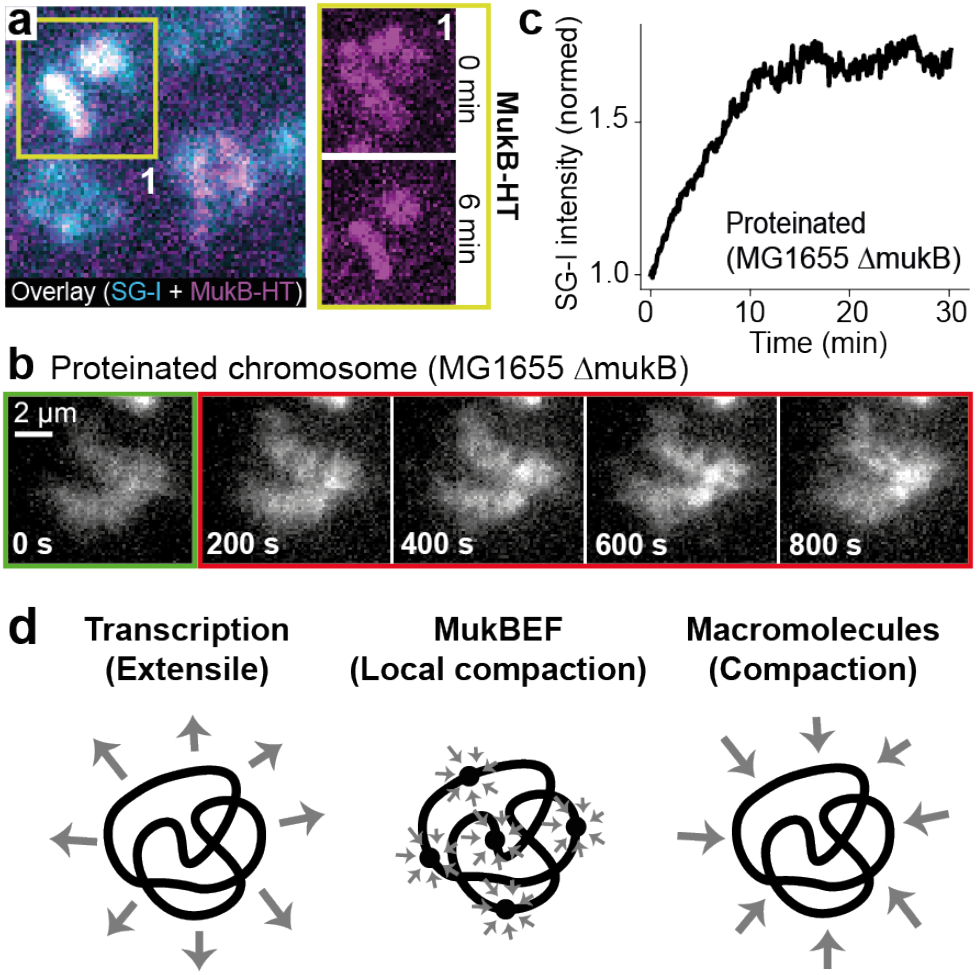
Dynamics of the loop extruder MukB on a cell-free chromosome. **a)** Representative fluorescence images of proteinated chromosomes with MukB-HT during cell-free gene expression. The chromosomes were labeled with SYBR Green I (SG-I) as intercalating DNA dye. A zoom on inset 1 shows a clustered distribution of MukB-HT on the chromosome after lysis and a more compact organization after 6 minutes of cell-free gene expression. The outcome was reproduced in two independent experiments. **b)** A fluorescence time-lapse montage with a zoom on a proteinated chromosome extracted from *E. coli* MG1655 Δ*mukB* transplanted into a large compartment. The cell lysate with active transcription (green box) was replaced at <200s by cell lysate without active transcription using rifampicin (red box). **c)** The average fluorescence signal of SYBR Green I (SG-I) was measured on five chromosomes extracted from *E. coli* MG1655 Δ*mukB* after transcription inhibition in a cell-free expression system (n=2 independent experiments). **d)** A simplified schematic of three processes acting on the chromosomes: Transcription as a source for genome-scale swelling, point-like compaction by local MukBEF clusters, and macromolecular crowding as a global compacting force.

### Gene expression dynamics on a cell-free chromosome

The *E. coli* genome has ∼700 α_70_ promoters, each regulating the production of mRNA with an approximate length of 1000 nucleotides, amounting to ∼15% of the total chromosome length (48). We sought to measure the cell-free transcriptional dynamics that induced chromosome swelling. However, a read-out through fluorescence labeling of all RNAPs in cell lysate would mask their dynamics on the chromosome because of their high bulk concentration. Instead, we used the fluorescently labeled RNAP and ribosomes bound to the proteinated chromosome from the source bacteria. As seen in Fig. 3, the RNAP signal again directly overlayed the HUα-GFP signal after lysis into cell-like compartments, presumably all bound to the chromosome (**Fig. 5a**). In comparison, the ribosome signal was shifted relative to the HUα-GFP signal (**Fig. 5b**), likely also bound to the fractured cell membrane by the translocation of membrane proteins (49). We introduced the cell-free gene expression system, tracking the expression dynamics through RNAP. The RNAP signal decayed evenly from the compartment within minutes, much quicker than the HUα-GFP decay, suggesting a fast evacuation of RNAPs from the cell-like compartment (**Fig. 5c, d**). We repeated the experiment in the presence of rifampicin to discern if the native RNAPs dissociated from the DNA after transcribing their genes or competed away by the incoming RNAPs from the cell-free gene expression system. Rifampicin did not change the decay in native RNAPs signal, excluding competition effects and suggesting native RNAPs finishing their transcripts on the chromosome. To gain further support, we omitted chemical energy from the reaction to observe a slower release of native RNAPs from the chromosome (**Fig. 5e**). We concluded that the genome-scale turnover of native RNAPs was homogeneous across the chromosome and occurred in less than 5 minutes.

**Figure 5:**
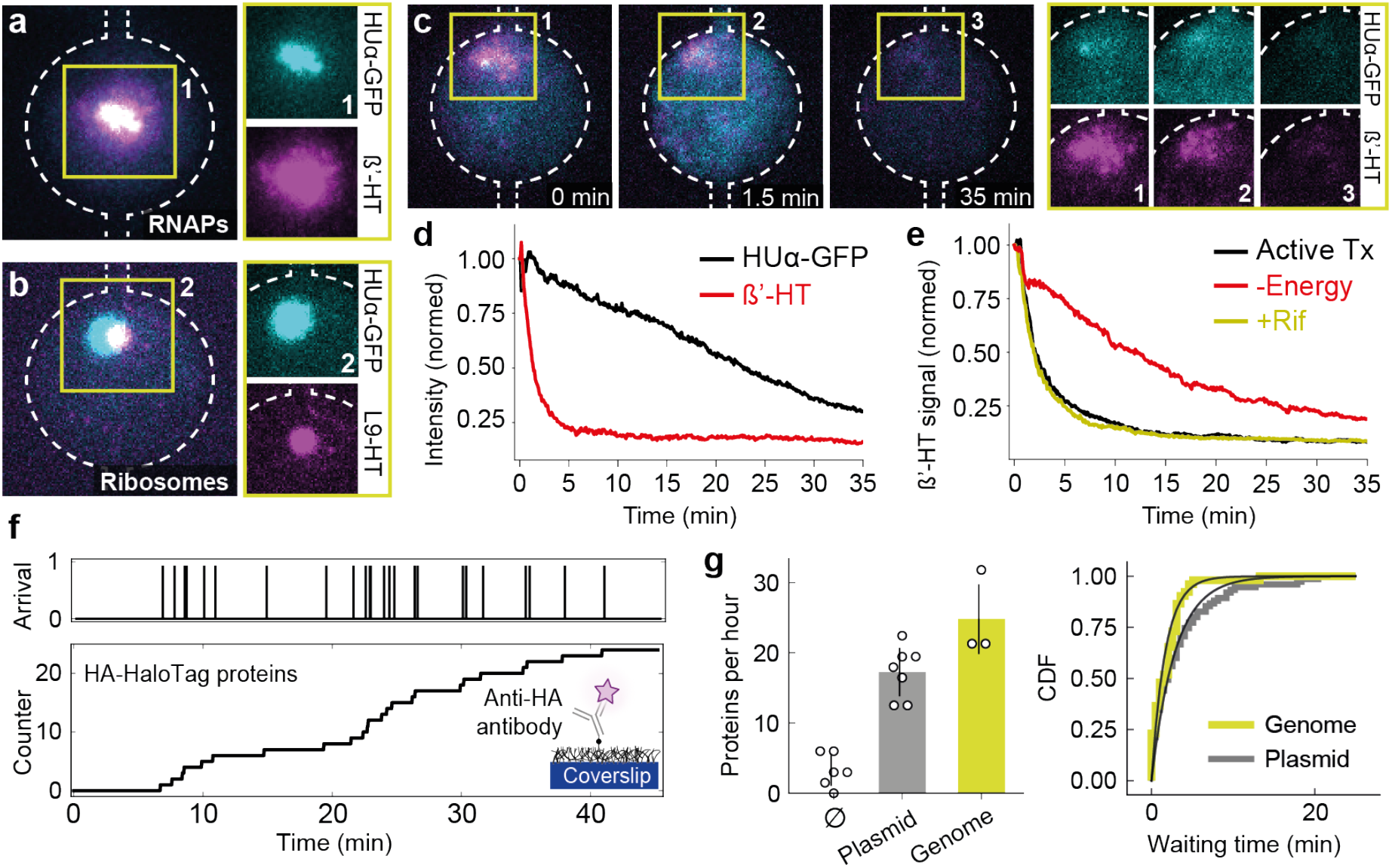
Cell-free gene expression dynamics with a proteinated chromosome. **a)** Representative double-color fluorescence image with embedded proteinated chromosomes. HUα-GFP was produced from plasmids, and β’-HT was produced from its native locus on the chromosome. The dashed white line outlines the cell-like compartment and capillaries connected with two main flow channels. **b)** A representative double-color fluorescence image of the embedded proteinated chromosome shows HUα-GFP produced from plasmids and L9-HT produced from the native gene locus on the chromosome. The dashed white line outlines the compartment and capillaries. **c)** Time-lapse fluorescence montage of the cell-like compartment loaded with a proteinated chromosome. HUα-GFP and β’-HT are shown in cyan and magenta, respectively. The dashed white lines outline compartments and capillaries. **d)** Fluorescence signals of HUα-GFP and β’-HT averaged over the compartment after introducing the cell lysate with active transcription. **e)** Fluorescence signal of β’-HT averaged over the compartment after introducing the cell lysate with active transcription (black curve), without energy for transcription (red curve), and with 500 nM rifampicin (Rif) to inhibit transcription initiation of the incoming RNAPs in cell lysate (yellow curve). **f)** An illustrative example of arrival times and integrated counts of HA-tagged HaloTag protein (purple star) during cell-free gene expression from a chromosome in cell-like compartments and binding onto anti-HA antibodies immobilized on the coverslip. **g)** The number of HaloTag spots detected during cell-free gene expression in cell-like compartments without DNA and loaded with single plasmids and chromosomes. The small circles show independent replicates, and the bars with error bars (S.D.) represent their means. The cumulative distributions of protein arrival times are shown with fits to mono-exponential distributions (black lines). The data was combined from independent experiments with plasmids (n=7) and proteinated *E. coli* chromosomes (n=3). All cell-like compartment diameters, 20 μm.

Intrigued by the turnover of RNAP from the chromosome, we next sought to measure protein synthesis as the output of the gene expression activity. We probed the typical activity of a gene by integrating a cassette on the chromosome with a native *E. coli* promoter, ribosomal binding site, and HA-tagged HT protein (HA: Human influenza hemagglutinin tag). The HA-tag allowed the capture of synthesized HT proteins on the surface of anti-HA antibody functionalized cell-like compartments for imaging (**Fig. 5f, Supplementary Fig. 7**) (38,50,51). Recording single-molecule HT appearances, we found production rates of ∼20 proteins per hour for the chromosome and, as a deproteinated reference commonly used in cell-free gene expression experiments, from single shorter recombinant DNA molecules with the same cassette (Methods, **Fig. 5g, Supplementary Fig. 7 and 8**). The similar protein synthesis from different DNA sources and our optical RNAP mapping (**Fig. 3e**) suggested that gene expression in our microfluidic compartments was independent of the chromosomes’ proteinated organizational state. Protein synthesis occurred at an approximate rate of one protein every three minutes, approaching an elongation rate of 6 nucleotides per second during transcription of the ∼1000-basepair long HT gene (and with the assumption that one mRNA produced one protein; no translational amplification).

With these numbers and the negligible amounts of MukB (for local compaction) in cell-free expression systems (Methods, **Supplementary Fig. 9a, b**) (52), we can estimate the free energy of transcription required to swell the deproteinated chromosome against macromolecular crowding. The integrated free energy from crowding should be around 10^5^ k_B_T (53), considering the osmotic pressure of a bacterial cytoplasm (6) and its ∼10-fold dilution during our cell lysate preparations. Each step during transcription dissipates a nucleotide with ∼44% of its chemical energy directed toward mechanical force (17,42). A single nucleotide holds 20 k_B_T, directing 8 k_B_T into motion, requiring 12,500 transcriptional steps to counterbalance crowding. If all ∼700 α_70_ regulated genes are covered by at least one RNAP, the 12,500 transcriptional steps are readily performed in one second, considering a transcription elongation rate of ∼10 nucleotides per second (like our estimate of ∼6 nucleotides per second) (54). Still, the calculation simplifies aspects such as gene direction and mechanical crosstalk through supercoiling. But without losing the expression activity, we could not directly test the response of DNA swelling by transcription to different crowding conditions (55,56).

### DNA displacement fluctuations under cell-free gene expression

Instead of different crowding conditions, we sought to quantify the DNA dynamics underlying the swelling and investigated structural changes in the deproteinated chromosomes during transcriptional activities. The DNA displacement fluctuations on the scale of a few genes were difficult to track because of the high local DNA density. We could only focus on high-intensity regions in the SG-I labeled chromosome as a proxy for semi-conserved structures (**Supplementary Fig. 9c**). With a histogram of jump distances and the mean-square displacements, we found low apparent diffusion coefficients of 10^−2^ - 10^−3^ μm^2^ s^-1^ with a subdiffusive scaling for deproteinated chromosomes (**Supplementary Fig. 9d, e**), very similar to diffusion coefficients found on isolated chromosomes in buffer conditions (57). Also, the displacement jumps lacked temporal correlation and together indicated an “active” diffusive behavior (**Supplementary Fig. 9f**) as observed on chromosomal loci in live cells (58).

The coarse tracking on the chromosome motivated us to devise an improved assay that could resolve DNA segments at the level of a few genes. To this end, we used shorter 48.5 kbps-long DNA derived from the lambda bacteriophage as a simplified testbed with only a few *E. coli* promoters (59). We labeled the DNA molecule at specific sites with fluorophores for imaging and with biotins for surface immobilization in open microfluidic chips (Methods, **Fig. 6a**). In addition to allowing a high-resolution tracking of the fluorescent spots, the site-specific labeling obviated the use of the intercalating SG-I dye, which might have slightly altered the DNA structure (60) and the transcription activities. We incubated the lambda DNA for about 30 minutes in independent experiments with cell lysate with and without active transcription and a dilute buffer lacking any proteins as control. After the incubation step, we started imaging the DNA molecules during a moderate flow pulse (∼40 seconds) along the channel, tracking the fluorescent DNA spots before, during, and after stretching by flow (**Fig. 6b**). We found no difference in the populations of DNA extensions between buffer and gene expression with transcription. Only the experiment with cell lysate and inhibited transcription reduced the average extension length when briefly stretched by flow (**Fig. 6c**). These findings suggest that the drag force could only stretch the swollen DNA molecule in the buffer and cell lysate with active transcription. Based on classic polymer physics (61), the swelling would further indicate that transcription effectively improves the solvent quality for DNA in cell lysate from poor to ideal or good.

**Figure 6:**
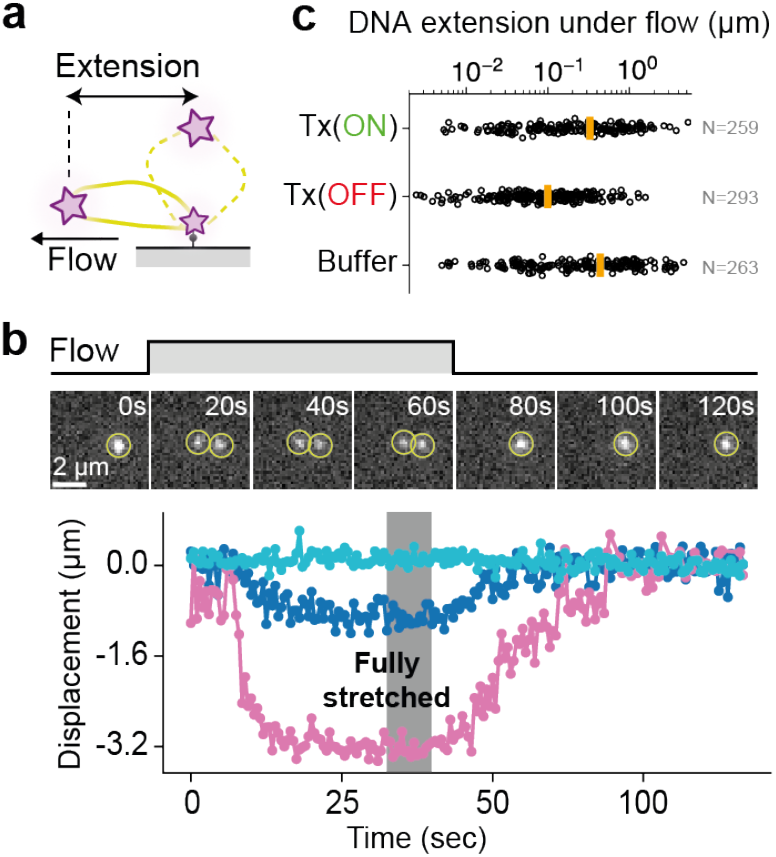
Transcription allows the stretching of shorter DNA molecules. **a)** A DNA molecule of the lambda bacteriophage was labeled on specific sites with fluorophores (dC-Cy5, purple star) for imaging and with biotins for surface immobilization in an open microfluidic chip. The surface-immobilized lambda DNA molecule (yellow dashed line) was stretched by flow (yellow solid-line circle). **b)** A representative fluorescence time-lapse montage shows a DNA molecule before, during, and after a moderate flow pulse. We tracked the position of the fluorescent dye to measure DNA extension, as shown for three different displacement examples. **c)** The DNA extension by flow was measured after incubating the DNA molecules for 30 minutes in cell-free expression systems with active transcription (Tx), without Tx by adding 500 nM rifampicin, and in phosphate-buffered saline (PBS) solution as a protein-free reference. The data were combined for Tx(ON), Tx(OFF), and buffer from n=3,2,3 independent experiments with mean values as yellow bars and individual molecules as circles.

## Discussion

The compacting and extensile behavior of the chromosome observed here is linked to the energy-consuming molecular machines operating on the DNA, rendering the system inherently out of equilibrium. This suggests that chromosomes are in the class of active gels (62), as soft materials with detailed balance locally broken. Together with polymer simulations and the recent technological achievements in assembling long synthetic DNA molecules (4,5,63–65), we will better understand the role of transcription for chromosome swelling on a molecular level in the future. The molecular driving forces could originate from various sources, including hydrodynamic flows (66,67), charge and steric effects by the nascent mRNAs, or transient DNA melting through transcription. Here, we focused on describing the fundamental effects on a native bacterial chromosome, which is likely important for all living and future artificial life-like systems.

## Acknowledgments

We thank L. Tunik of the Nanofabrication unit at the Weizmann Institute for support in the manufacturing process, the Forchheimer plasmid collection for the bacterial strain and plasmids, D. Garenne for helping prepare the cell lysate, Y. Barak for helping with the strain preparations, and A. Kumar and D. Bensimon for critically reading the manuscript. The K-12 MG1655 rpoC-HT, L9-HT, and both mukB-HT and ΔmukB strains were a generous gift from J. Mäkelä, M. Johansson, and F. Bürmann, respectively.

## Author contributions

F.G. conceived of the project, performed experiments, analyzed experiments, drafted and edited the manuscript; S.S.D. discussed the data, drafted and edited the manuscript; R.B.Z. discussed the data, drafted and edited the manuscript; V.N. discussed the data, drafted and edited the manuscript.

## Data availability

Data supporting the findings of this study are available in the article and its Supplementary Information section. Raw fluorescence movies will be deposited on Zenodo.

## Supplementary Materials

Figs. S1 to S9 Tables S1 and S2

References

## Methods

### Design, fabrication, and preparation of microfluidic chips

We designed the microfluidic chip layout with AutoCAD 2021 (Autodesk) and exposed 5” chrome masks (Nanofilm) with the MicroWriter ML 3 (Durham Magneto Optics Ltd). The mask was developed according to the manufacturer’s protocol. Next, a 4” silicon wafer (0.525 mm thickness, <100>, p-type, University Wafers) was cleaned with acetone, isopropanol, and water. The wafer was cleaned with a plasma system (250 W, O_2_ at 40 sccm, 150 mTorr, 120 sec; March AP-300, Nordson). To promote adhesion of the SU-8 mold, the wafer surface was prepared with Hexamethyldisilazane (HMDS, Transene Company) for 30 s (after incubation, the wafer was dried by spinning for 30 sec at 3000 rpm), and a base layer from SU8 2000.1: 1) 7.5 sec at 500 rpm, 2) 30 sec at 1500 rpm with global UV exposure and hard bake. The first mold layer was SU-8 2000.5. The resist was spun on the wafer: 1) 7.5 sec at 500 rpm, 2) 30 sec at 1900 rpm. The resist was exposed with a mask and mask aligner (Karl Suss Ma6/BA6), baked, and developed using the manufacturer’s protocol. The second layer was SU-8 6001. The resist was spun on the wafer 1) 7.5 sec at 500 rpm and 2) 30 sec at 1500 rpm. The resist was aligned with the first layer, exposed, baked, and developed. The final layer was fabricated with SU-8 3050: 1) 7.5 sec at 300 rpm, 2) 30 sec at 2000 rpm. The resist was first exposed after aligning the first and second layers and then baked and developed. For the plasmid experiments, the SU8 mold was constructed from only two layers where the compartment and both capillaries were made from SU8 2000.5 and the flow channel from SU-8 3050. After verifying the SU-8 structures, the resist was hard-baked at 150 °C and slowly cooled to room temperature. The final dimensions were measured with a profilometer (Dektak, Bruker) and optical microscopy.

The wafer was placed in a petri dish and covered with ∼40 mL of polydimethylsiloxane (10:1 PDMS:Curing agent, Sylgard 184, Dow Corning). The dish was placed for ∼1 h under vacuum to remove bubbles and baked at 75-80 °C for ∼4 h. The PDMS was cut into pieces using a scalpel and peeled off the wafer. Holes for the inlet and outlet were punched (0.75 mm diameter, Welltech Labs) on a cutting mat. The microfluidic chips were cleaned with isopropanol and blow-dried. The coverslips (#1.5, Marienfeld) were cleaned by boiling in 1) 96% ethanol for 10 min and 70 °C warm and 2) 3:1:1 H_2_O:NH_3_(25%):H_2_O_2_. The coverslips were blow-dried and stored in a dust-free box. Before bonding PDMS to the coverslip, we again cleaned the coverslips with the plasma system (250 W, O_2_ at 40 sccm, 150 mTorr, 120 sec). Then, the PDMS and coverslips were surface activated (100 W, O_2_ at 49 sccm, 180 mTorr, 20 sec) and brought in close contact for permanent bonding. The assembled microfluidic chips were baked at 75-80 °C for ∼4 h.

The microfluidic chips were incubated with ∼3 mg of photoactivatable silane polyethylene glycol (PEG) (68) in 1.5 mL of anhydrous acetonitrile (Sigma-Aldrich) for ∼30 min. After incubation, the microfluidic chips were successively washed with acetonitrile, an equimolar mix of acetonitrile and water, and water. The photo-sensitive PEG biochip was exposed with a UV cube (UV KUB), treated with 5 mg ml^-1^ NHS-Biotin (ThermoFisher Scientific) in 250 mM borate buffer (pH 8.6) for 20 min, and flushed with degassed 0.1% Tween-20 in PBS (phosphate-buffered saline, pH 7.4). The microfluidic chips were kept wet in a humid chamber until use (typically for 0-2 days).

### Defining the dimensions of cell-like compartments in microfluidic chips

The lifetime of molecules in the compartment is quantified by the layout and diffusion coefficient *D* of the molecule (35)

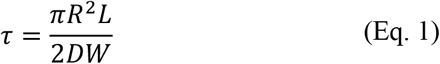

with *R, L*, and *W* as the compartment’s radius, and the capillary’s length and width, respectively. To optimize the compartment layout, we initially constructed cell-like compartments with R=5 μm, L=40 μm, and W=1.5 μm for single-molecule tracking experiments. We contained a single plasmid, tracked the plasmid position (pBEST HA-HT) with TrackMate (69), and fitted the mean-square displacement to normal diffusion ⟨*r*^2^⟩ = 4*Dt* (**Supplementary Fig. 8**). We found a typical diffusion coefficient of D_D_ ∼ 1.7 μm^2^ sec^-1^. We then designed a layout with a larger compartment size (R=10 μm) to contain the *E. coli*chromosome for more than 1 hour and the plasmid for ∼40 min, long enough for our expression experiments. The layout further gave a lifetime for mRNA τ_M_ around 7 min with a typical diffusion coefficient of D_M_ ∼ 10 μm^2^ sec^-1^ (70), but was reduced to a few minutes (comparable to cells) through enzymatic degradation in cell lysate (35). The fast degradation presumably prevented mRNA from escaping the cell-like compartment. It allowed us to record the integrated translation activity throughout the mRNA’s entire lifetime in cell lysate by surface-capturing and imaging the nascent proteins.

### Experimental setup with single-molecule fluorescence microscope

Fluorescence images were acquired using Micro-Manager 2.0.0 and a custom-built single-molecule TIRF microscope, as previously described (38). The lasers with 488 nm (100 mW, OBIS, Coherent) and 647 nm (120 mW, OBIS, Coherent) excitation were combined (DMSP605 and 5xBB1-E02, Thorlabs) via an objective mounted on an XYZ stage (MBT612D/M, Thorlabs) into a single-mode fiber (P5-460B-PCAPC-1, Thorlabs). The fiber was coupled into a mirror collimator (RC08FC-P01, Thorlabs) to expand the laser diameter to 8 mm, guided through an achromatic lens (f = 150 mm, AC254-150-A-ML, Thorlabs), redirected by a mirror and filter cube (TRF59906, Chroma), and focused onto the back focal plane of the TIRF objective (60x, 1.49 NA, Nikon). The emission was focused with a tube lens (TTL200, Thorlabs) onto an EM-CCD (iXon Ultra 888, Andor Technology, Belfast, UK). The two lasers were controlled with an Arduino microcontroller (Arduino control software v1.2) and synchronized with the output trigger signal of the EM-CCD camera. The sample was translated in XY (Märzhäuser Wetzlar), and the TIRF objective was mounted with an in-house built Delrin adapter for thermal insulation on a piezo stage to focus in Z (400 μm Fast PIFOC, PI). The objective was further enclosed with resistive heating foil (HT10K, Thorlabs) to set the temperature at 34 °C using a PID controller (TE-48-20, TE Technology). We constructed a custom humidity chamber and flushed it with humid N_2_. We acquired widefield fluorescence images, but the refractive index mismatch between PDMS and glass would also allow total-internal reflection fluorescence (TIRF) microscopy. Solutions were pulled from a 10 μL pipette tip attached to the inlet port through the microfluidic chip with a glass syringe (Gastight, Hamilton) and syringe pump (PHD Ultra, Harvard Apparatus).

### Engineering the E. coli genome

We transformed *E. coli* K-12 MG1655 with the plasmid pSLTS (Forchheimer plasmid collection, Bacteriology and Genomic repository, Weizmann Institute) and induced the lambda red system for homolog recombination (71). To generate the donor plasmid pKD4-HT, we transferred the previously established HaloTag reporter system with N-terminal fused HA tag (**Supplementary Table 1**) (54) and the regulatory sites for transcription and translation from pBEST into pKD4 (72) (Forchheimer plasmid collection, Bacteriology and Genomic repository, Weizmann Institute) using PirPlus DH10β pir116 (Bacteriology and Genomic repository, Weizmann Institute) as cloning host. The plasmid was constructed by DNA assembly (NEBuilder HiFi DNA Assembly, NEB) of PCR fragments, amplified using KAPA HiFi HotStart ReadyMix (Kapa Biosystems, Roche), and verified through in-house sequencing.

The MG1655 strain with induced lambda red system was transformed with the PCR product amplified (**Supplementary Table 2**) from pKD4-HT containing the HA-HT expression cassette, Kanamycin selection marker, and homologous regions. Several colonies were picked from agar plates with Ampicillin and Kanamycin for colony PCR with two flanking primers to verify the chromosomal insertion (**Supplementary Table 2**). PCR fragments with correct fragment lengths were sequenced. The final strain MG1655 P_R_ UTR1 HA-HaloTag was then cured from pSLTS by growing the cells at 37 °C on agar plates with Kanamycin. Growth curves for all strains were measured in a 24-well plate (Greiner) using a plate reader (ClarioStar Plus, BMG Labtech). A volume of 1.5 mL lysogeny broth (LB) medium was inoculated with 15 μL overnight cultures (inoculated from -80 °C glycerol stocks) to measure the OD_600_ values every 3 minutes at 30 °C.

### Plasmid construction for HUα-GFP

The plasmid pBEST HUα-GFP was derived from P70a-UTR1-deGFP with Ampicillin resistance (73). We amplified the gene *hupA* from *E. coli* K-12 MG1655 with colony PCR (KAPA HiFi HotStart ReadyMix, Kapa Biosystems, Roche) and primers (IDT, **Supplementary Table 2**). We fused the gene to the N-terminus of the *degfp* gene, separated by a short peptide linker (KRAPGTS). We assembled the PCR fragments with the NEBuilder HiFi DNA Assembly (NEB). Next, we reduced the transcription activity with a version of the PLlacO1 promoter (**Supplementary Table 1**) with PCR and primers, circularized the linear PCR fragment with a KLD Enzyme Mix (NEB), cloned with chemically competent DH5α cells, verified the construct by sequencing, and transformed the final plasmid into the various strains.

### Extraction of E. coli chromosomes

To extract the chromosome from *E. coli* cells, we followed the protocol by Pelletier *et al. (6)*. We grew the strains overnight, transferred 100 μL into 10 mL fresh LB medium, and Ampicillin the next day to grow cells at 30 °C for ∼2 hours, reaching exponential phase (OD∼0.4). Because deleting *mukB* renders *E. coli* highly temperature-sensitive with defects in chromosome segregation (43), we grew only the MG1655 ΔmukB strain overnight at ∼22 °C to be directly processed without the second dilution and incubation step. We centrifuged 1.5 mL of cell suspension for 2.5 min at 5,000xg to harvest the cells. The supernatant was gently removed. The cells were washed in 500 μL PBS buffer and then resuspended in 500 μL of sucrose buffer (20 % sucrose (w/v), 100 mM NaCl, 10 mM EDTA, 10 mM NaPi, pH 7.3). The cells were centrifuged again and incubated with 2.5 μM MaP655-Halo (54) (stock stored at -20 °C in dimethyl sulfoxide, DMSO, Sigma) in 100 μL sucrose buffer for 1 hour, followed by centrifuging and washing in 500 μL sucrose buffer. The cells were gently mixed in a 2 mL tube with a rotisserie during incubation.

To test the labeling efficiency of the HaloTag with MaP655-Halo, we also performed a control experiment by blocking the HaloTag with HaloTag Biotin Ligand (Promega). Bacteria in 100 μL sucrose buffer were first incubated with 2.5 μM HaloTag Biotin Ligand (stock stored at -20 °C in DMSO) or 1 μL DMSO (stored at -20 °C). After a wash step with 500 μL sucrose buffer, the cells were incubated with MaP655-Halo as described above and imaged without cell lysis. The HUα-GFP signal was used to segment bacteria trapped in large compartments to measure the HT signal with the MG1655 rpoC-HT strain as a highly and constantly expressed gene (**Supplementary Fig. 6**).

For on-chip cell lysis, the plasmolyzed cells were harvested by centrifuging, resuspended in 75 μL sucrose buffer, flushed onto the chip already filled with sucrose buffer, and introduced into the compartments by centrifuging the microfluidic chip mounted on a tilted stage at a speed of 500×*g* for 10 minutes. Next, we introduced 300 μg mL^-1^ lysozyme from chicken egg white (Sigma-Aldrich) in sucrose buffer and incubated the chip for 1 h at room temperature to degrade the cell wall. We changed the buffer on the microscope-mounted chip at a flow rate of 1.5 μL min^-1^ with a hypotonic solution (100 mM NaCl, 0.5 mg ml^-1^ BSA, 20 mM Hepes, pH 7.5) to lyse the spherical cells. Chromosome-bound proteins were degraded by flushing hypotonic solution supplemented with proteinase K (PKA, stock concentration of 800 units mL^-1^, NEB). The mix was prepared by adding 0.75 μL PKA to 20 μL hypotonic solution.

### Local pulling of proteinated chromosomes with an electric field

After cell lysis, a ∼5 V cm^-1^ electric field was manually applied across the compartment using a DC power supply (E3620A, Agilent) to pull the chromosome into the bottom capillary. Electrodes were constructed from standard electric wires and installed into a second pair of microfluidic in- and outlet ports. The fluorescence profile along the capillary was measured using Fiji v1.0 and plotted as a kymograph using the command “Reslice”. The kymograph was loaded in Python v3.7 and processed using scikit-image v0.17.2 and NumPy v1.20.3.

### Preparation of Escherichia coli-based cell-free gene expression systems

The cell lysate was prepared with *Escherichia coli* (BL21 Rosetta 2) and K-12 MG1655 strains (73). Briefly, bacteria were grown in 2xYT supplemented with phosphates. The bacteria were lysed after reaching an OD600 = 1.5–2 using a pressure cell. After the first centrifugation step (12,000×*g* for 10 min), the supernatant was incubated at 37 °C for 80 min for a run-off reaction. During the run-off reaction, we optionally added the HaloTag immobilization beads (Magne HaloTag beads, Promega) to pull down MukB-HT, providing a background reference to estimate the MukB-HT concentration in the cell lysate without pull-down. A bead volume of 100 μL was washed twice with S30B (14 mM Mg-glutamate, 60 mM K-glutamate, 5 mM Tris, pH 8.2) buffer following the manufacturer’s protocol, added to the run-off reaction, and later removed with a magnet. After a second centrifugation step at 12,000×*g* for 10 min, the cell lysate’s supernatant was dialyzed for 3 hours at 4 °C. After a final spin-down at 12,000×*g* for 10 min, the supernatant was aliquoted (29 μL) and stored at −80 °C. Before use, the cell lysate was thawed on ice, supplemented with the necessary solutions (10 mM Mg-glutamate, 80 mM K-glutamate, 4% PEG8000, 10.8 mg/ml Maltodextrin, amino acids, energy buffer, and nuclease inhibitor GamS), filled to 78.3 μL with water, and gently mixed with a pipette. Finally, 9 μL of cell lysate was mixed with 1 μL of water containing reagents such as plasmid or fluorogenic dye.

### Performing chromosome expression experiments in large and cell-like compartments

Cells were lysed for 15 min under a flow of 1.5 μL min^-1^. The lysis buffer was exchanged under a flow of 0.5 μL min^-1^ by a cell-free gene expression system supplemented with 100 nM SYBR Green I (Thermo Fisher Scientific) (60) for experiments in the large compartments, and, in the case of inhibited transcription, 500 nM rifampicin (15 μM stock solution was stored in DMSO at -20 °C). We omitted the energy buffer in preparing the cell-free expression system to check for reduced transcription activity on the proteinated chromosome in the cell-like compartments. Fluorescence images were acquired with an excitation power of 3 W cm^-2^ (488 nm) and 15 W cm^-2^ (647 nm), 20 ms exposure time, and 750 gain. Images were acquired every 2.5 seconds for alternating double-color imaging and every 5 seconds for single-color imaging.

### Image segmentation of chromosomes in large compartments

Cell-free chromosomes were segmented using StarDist v0.9.1 (74) and tracked using TrackPy v0.6.4 to measure the SYBR Green I signal and estimate the chromosome area. We used the pre-trained StarDist model “2d_versatile_fluo” for 2D to segment chromosomes in the large diamond-shaped compartments. The segmented chromosomes were linked into trajectories when the weighted centroids were within a search range of 0.8 μm and returned after vanishing for a maximum of 3 frames. The segmentation and tracking results were manually verified using Napari v0.5.1.

To find the fraction of MukB-HT spots on DNA blobs during the cell-free gene expression experiments, we localized and tracked MukB-HT spots and SG-I labeled DNA blobs on the chromosome using custom-written software (https://github.com/FerdinandGreiss/nanokit). We deemed the two spots colocalized when MukB-HT and DNA blobs were closer than 0.32 μm (or 2 pixels).

### Functionalization of cell-like compartments with antibodies

We followed the steps described by Vonshak *et al*. (50). To functionalize the cell-like compartments with anti-HA antibodies, we mixed the biotinylated high-affinity (3F10) anti-HA antibodies (50 μg ml^−1^, ∼500 nM, Sigma-Aldrich) at a 2:1 ratio with Streptavidin in PBS. We incubated the solution on ice for ∼30 min. The antibodies conjugated with Streptavidin were incubated on the chip. After ∼20 min, the main flow channel was thoroughly flushed with PBS to remove unbound antibodies.

### Measuring protein synthesis from chromosomes in cell-like compartments

One hour after cell lysis and the transplantation of the chromosome into cell-like compartments, the solution in the microfluidic chip was carefully changed to PBS. The long waiting time allowed freely diffusing HT proteins to evacuate the cell-like compartments. The compartments were then functionalized with anti-HA antibodies, as described above. After carefully removing the unbound antibodies with PBS, we introduced cell lysate and fluorogenic dye MaP655-Halo at a final concentration of 50 nM. We picked single cell-like compartments with intense HUα-GFP signals and acquired fluorescence images with an excitation power of 3 W cm^-2^ (488 nm) and 15 W cm^-2^ (647 nm) at a frame rate of 1 Hz (in alternating mode between the two wavelengths; the time delay between frames of a single excitation wavelength is therefore 2 secs), 100 ms exposure time, and 750 gain.

The fluorescent single-molecule spots were identified, localized, and tracked using custom software (https://github.com/FerdinandGreiss/nanokit). Furthermore, as only a few proteins were produced during an experiment, we linked protein spots with a clustering algorithm (scikit-learn v0.24.2, dbscan function) to reduce overcounting due to the temporal cut-off during the tracking process. Also, spurious spots outside cell-like compartments were removed from the analysis by manually outlining the cell-like compartment.

### Performing single plasmid expression experiments in cell-like compartments

The plasmid encoding the HA-HT gene was fluorescently labeled using a nick-translation protocol (54). We labeled the plasmids with Atto647N as a far-red fluorophore to minimize the autofluorescence background from the cell lysate. Despite an overlap between the signals of the fluorogenic MaP655-Halo dye and plasmid, misidentification between protein and DNA was minimal due to the fast bleaching time of DNA, on the order of a few minutes (54). After incubating the antibody-functionalized microfluidic chip (prepared as described above) with 1 nM of fluorescently labeled plasmids for ∼1 min, 10 μL of cell lysate and 50 nM fluorogenic dye were introduced to remove residual plasmids in the main flow channel and to initiate gene expression in the cell-like compartments. We picked compartments containing a single diffusing DNA molecule and started imaging. Fluorescence images were acquired with an excitation power of 15 W cm^-2^ (647 nm) and at a frame rate of 1 Hz, 100 ms exposure time, and 750 gain.

### Cell-free gene expression with surface-immobilized lambda DNA

2500 ng lambda DNA (NEB) was mixed with 1x Cutsmart Buffer (NEB), 1 μM dG, dG, dT, dA-Biotin (Jena Biosciences), 5 units DNA Pol I (NEB) with water in a total volume of 2x 20 μL. The mix was incubated for 1 hour at ∼22 °C. The solution was cleaned using magnetic beads (Sera-Mag Select, Cytiva) according to the manufacturer’s protocol and eluted with 15 μL. The purified lambda DNA was further nick-translated with 25 μM dG, 25 μM dA, 25 μM dT, 15 μM dC, 10 μM dC-Cy5 (Jena Biosciences) and cleaned as described above. The modified lambda DNA was then ligated in 20 μL reaction volume with 1x Cutsmart buffer, 1 mM ATP, and 5 units T4 DNA Ligase (Thermo). The mix was incubated at 16 °C for 2 hours, heat-inactivated by incubation at 65 °C for 10 minutes, and further nicked with 5 units Nb.BbvCI (NEB) at 37 °C for 1 hour, followed by a heat-inactivation step at 80 °C for 20 minutes. The nicks were translated by adding 25 μM dG, 25 μM dA, 25 μM dT, 15 μM dC, 10 μM dC-Cy5, together with 5 units DNA Pol I (NEB). The mix was incubated for 1 hour at ∼22 °C and cleaned using magnetic beads. The final DNA preparation with a typical concentration of 40 ng μL^-1^ (∼1 nM) was stored at -20 °C.

The modified lambda DNA was incubated with Streptavidin (Sigma-Aldrich) in PBS at a final concentration of 180 pM and 4 nM, respectively. We used the same DNA preparation for all three experimental conditions to remove any potential issues with batch-to-batch variations in labeling and breakage of the long lambda DNA during the preparation steps. The Streptavidin-conjugated DNA was incubated for 10 minutes in an open microfluidic chip prepared as described above. The chip had 5 thinner parallel flow lanes, each with a height of 70 μm and width of 200 μm. The parallel microfluidic flow channels allowed us to measure the different conditions on the same chip. The chip was washed with PBS to remove residual DNA, flushed with a cell-free gene expression system with and without 500 nM rifampicin, and PBS as a reference for a 30-minute incubation at 34 °C. After incubation, we acquired ∼10 images, started the syringe pump with a moderate flow rate of 2.25 μL min^-1^ to stretch the surface-immobilized DNA for the next 75 frames, and stopped the flow to record the DNA molecules relaxing back into their original configurations for another ∼100 frames. The flow rate was not enough to resolve all seven Nb.BbvCI nicking sites on the lambda DNA. During the stretching process, images were acquired every 0.5 seconds with an exposure time of 100 ms, 750 EM Gain, and 15 W cm^-2^ (647 nm). We analyzed the images by identifying and tracking the fluorescent spots (https://github.com/FerdinandGreiss/nanokit) to extract the average positions between frames 65 and 75 (stretched) and after frame 150 (relaxed). The displacement between these two positions gave the DNA extensions by flow.

