## Supplementary Information for "Transcription-dependent swelling of a transplanted chromosome in an artificial cell"

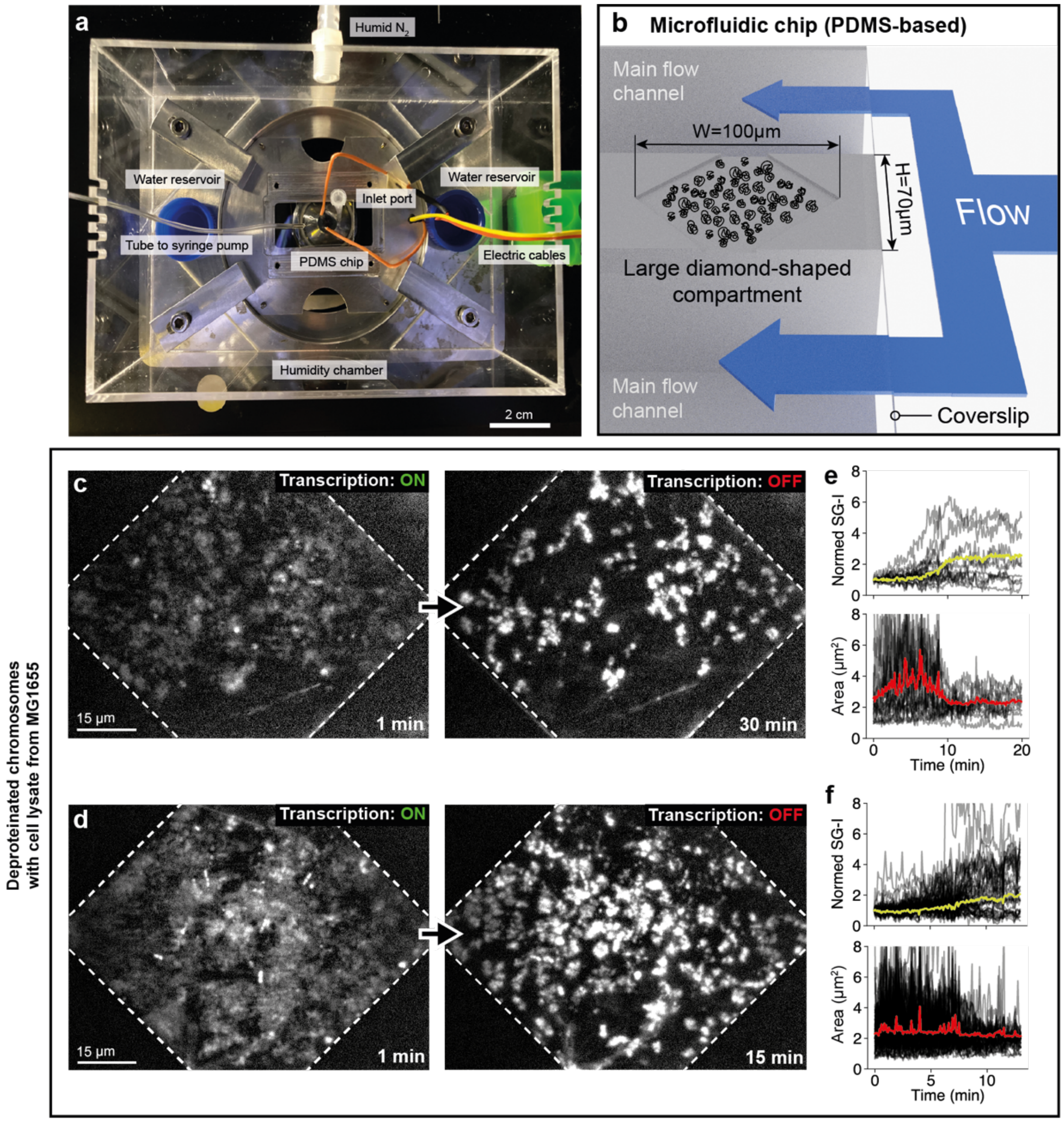


**Supplementary Figure 1: The experimental system interfaced with electronics, fluidics, and a single-molecule fluorescence microscope and compaction of cell-free chromosomes.**

**a)** The experimental setup with a polydimethylsiloxane-based microfluidic chip integrated into a single-molecule fluorescence microscope system, connected to electronics, and stabilized against environmental fluctuations through a custom humidity chamber. **b)** A 3-D rendering of a diamond-shaped compartment for steady-state experiments, loaded with cell-free chromosomes and flanked by two main flow channels for the fluid exchange. **c)** Fluorescence images (before and after transcription inhibition) of deproteinated chromosomes in the diamond-shaped compartments with lysate extracted from K-12 MG1655. **d)** Another example of deproteinated chromosomes transitioning from active to inactive transcription in an MG1655-based cell-free expression system. **e)** Analysis of panel c: The shift from active to inactive transcription in the large compartment was marked by a ~50% drop in estimated chromosome area and a ~2.7-fold increase in the SYBR Green I (SG-I) signal. Identified chromosomes are plotted as black curves, their median as red (area) and yellow (normed SG-I signal) curves. **f)** SG-I intensity and area analysis of panel d. Identified chromosomes are plotted as black curves, their median as red (area) and yellow (normed SG-I signal) curves.

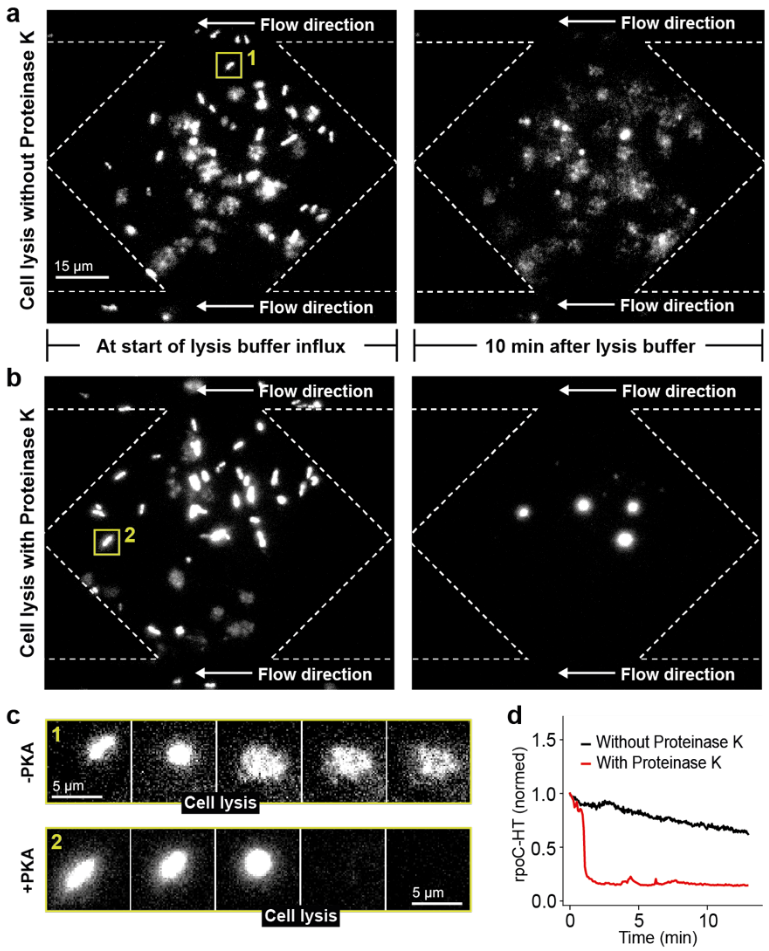


**Supplementary Figure 2: Cell lysis and protein degradation in large steady-state compartments.**
**a)** Representative fluorescence images of *E. coli* MG1655 (labeled with β'-HT) at the beginning of cell lysis and 10 minutes later without Proteinase K (PKA). **b)** Representative fluorescence images of *E. coli* K-12 MG1655 (labeled with β'-HT) at the beginning of cell lysis and after 10 mins of cell lysis in the presence of PKA. **c)** Two fluorescence time-lapse montages with zoom on single bacteria undergoing lysis without (upper row) and with (lower row) PKA in lysis buffer. The cells are indicated by yellow boxes in panels a and b. **d)** Normalized fluorescence intensity traces of a single bacterium labeled with β'-HT. PKA led to the fast decay in the β'-HT signal after cell lysis, suggesting a fast degradation of all proteins attached to the chromosome. Lysis without PKA maintains the chromosome-bound β'-HT.

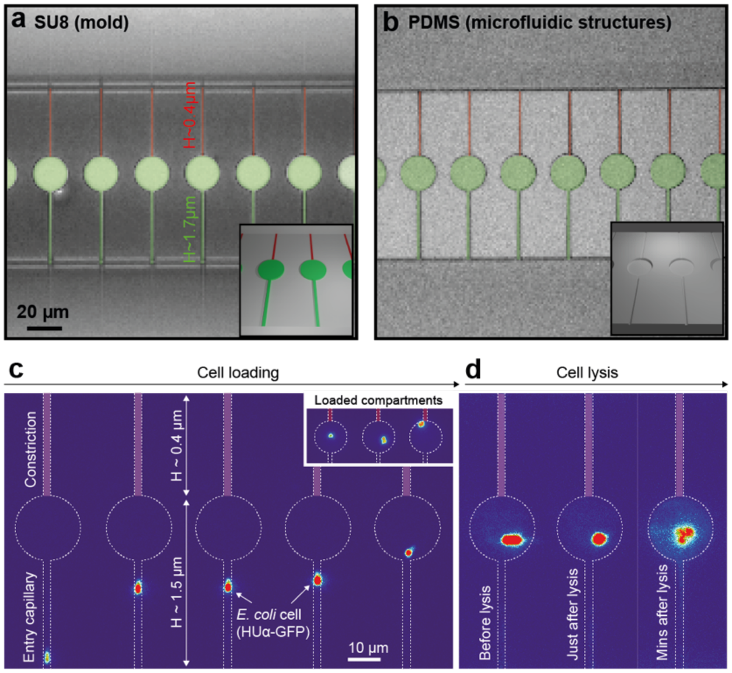


**Supplementary Figure 3: Cell-like compartments to study individual cell-free chromosomes.**
**a)** 3-layer SU8 mold with the first layer (colored in red) at a height of ~0.4 µm, the second layer (colored in green) at a height of ~1.5 µm, and the third layer at a height of ~75 µm. The third layer produced the main flow channels. **b)** A brightfield image of the polydimethylsiloxane (PDMS) chip after baking the uncured elastomer on the SU8 mold. **c)** The cell-like compartments are loaded with single *E. coli* bacteria through centrifugation, exemplified by a time-lapse series of a HUα-GFP (plasmid) labeled bacterium entering a cell-like compartment. The inset shows three compartments loaded, each with a single bacterium. **d)** After loading the plasmolyzed cells, we degraded the cell wall on the chip and exchanged the buffers to lyse the cell in situ.

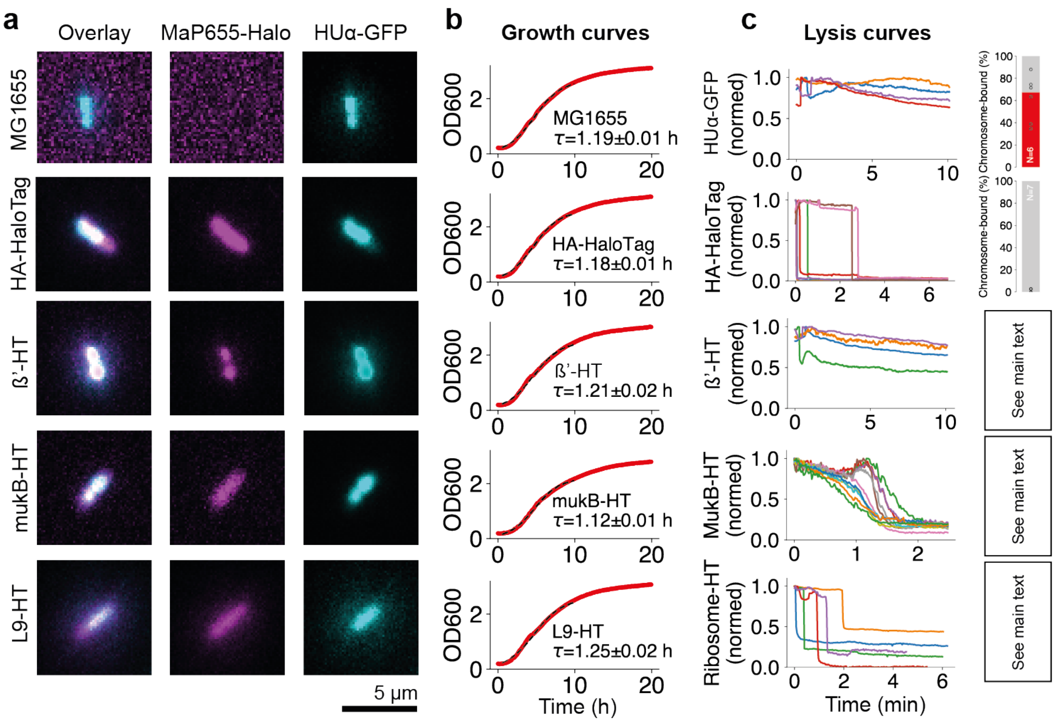


**Supplementary Figure 4: The collection of bacterial strains with genes tagged with HaloTag and their growth curves and lysis properties.**

**a)** All strains were derived from K-12 MG1655 with HUα-GFP expressed from plasmids and incubated with 2.5 µM fluorogenic dye MaP655-Halo before imaging. **b)** The growth curves were measured in a plate reader in lysogeny broth (LB) medium at 30 °C. The red curves were averaged from three replicates in a well plate and fitted to the logistic equation OD600 = $K/(1-(1-K/N_0)2^{-t/\tau}$ (shown as black dashed line), estimating the doubling times τ for each strain. **c)** The cell lysis curves were produced from single bacteria trapped in compartments. We tracked the fluorescence signal and computed the ratios before and after cell lysis to find the fraction of chromosome-bound proteins.

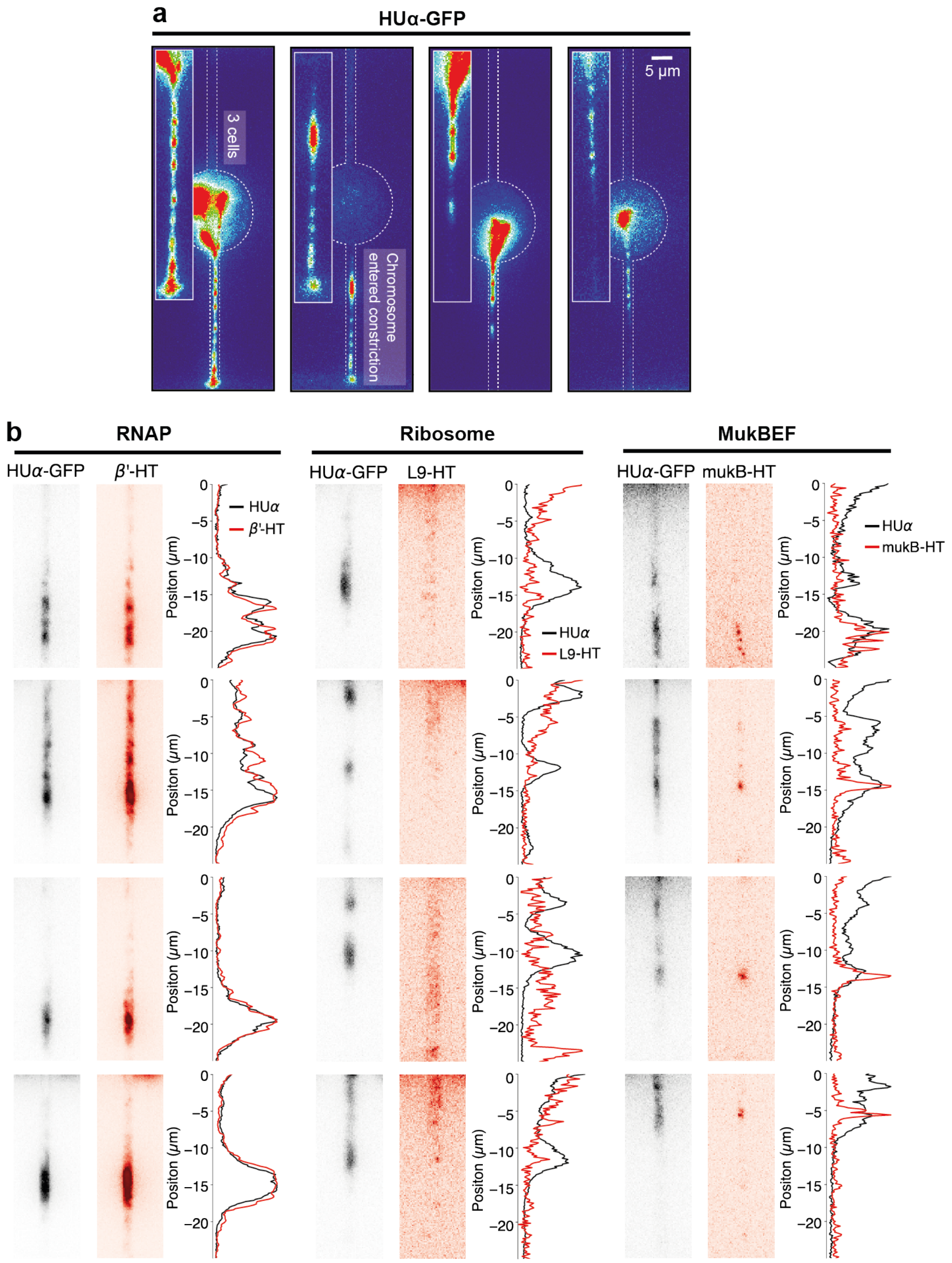


**Supplementary Figure 5: Individual *E. coli* chromosomes stretched by an electric field.**

**a)** The *E. coli* chromosomes were labeled with HUα-GFP, extracted *in situ*, and gently pulled into the lower capillary using a weak electric field. Here, four examples are shown, with one chromosome completely pulled into the small capillary (second panel from left) and another where three cells were lysed in a single compartment (leftmost panel). All cell-free chromosomes showed persistent bright HUα-GFP blobs along the stretched chromosome. **b)** Further four snapshot examples of stretched chromosomes with HT-labeled RNAP (β'-HT, left column), ribosome (L9-HT, center column), and MukBEF (MukB-HT, right column) in cell-like compartments using an E-field.

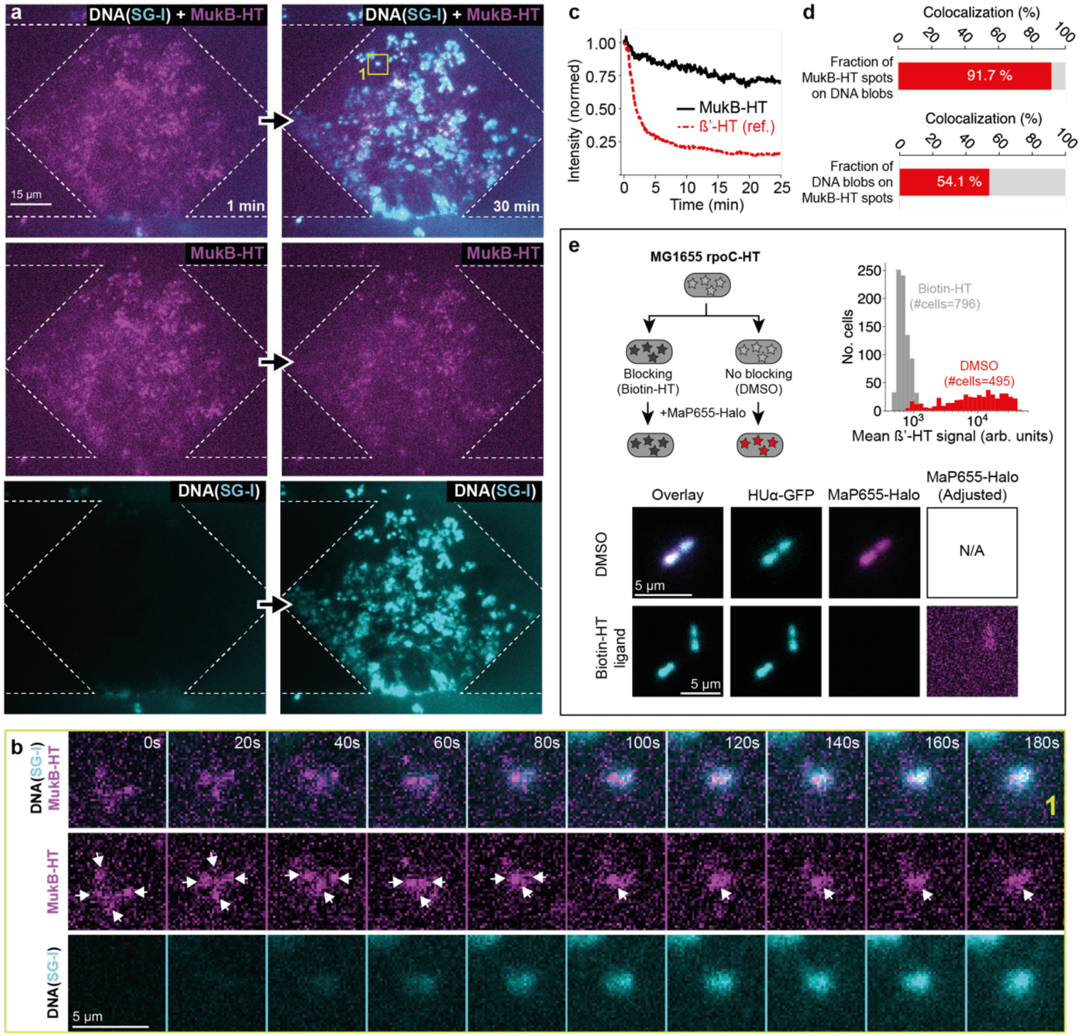


**Supplementary Figure 6: Proteinated cell-free chromosomes, cell-free mukB-HT dynamics, and HT labeling *in vivo*.**

**a)** Fluorescence images of a cell-free gene expression experiment with proteinated chromosomes extracted from MG1655 mukB-HT. MukB-HT was labeled before cell lysis with MaP655-Halo, showing clusters of MukB-HT on the cell-free chromosomes. The DNA intercalating dye SYBR Green I (SG-I) was flushed with the cell-free gene expression system into the microfluidic chip, labeling the DNA shortly after the start of the activity experiment. **b)** Fluorescence time-lapse montage with zoom on a chromosome marked with a yellow box in panel a. **c)** Fluorescence signals on single chromosomes trapped inside large compartments for β'-HT (reference) and MukB-HT. The decay curves were reproduced in two independent experiments. **d)** The MukB-HT spots localized with 91.7% on a DNA blob. In comparison, a DNA blob colocalized to a MukB-HT spot with 54.1% because of minor inhomogenous HT labeling (see panel e). **e)** Control HT-labeling assay in *E. coli* cells without lysis: Bacteria were first incubated with Biotin-HT (neg. control) and Dimethyl sulfoxide (DMSO). Both cell preparations were then incubated with the fluorogenic dye MaP655-Halo. The HT labeling was reduced to negligible signal levels when first blocked by the Biotin-HT ligand. Still, a minor non-labeled peak around $10^3$ arb. units also remained without blocking using DMSO. The HT signals were measured with MG1655 rpoC-HT bacteria as a highly and constantly expressed gene from two independent experiments to produce the histograms.

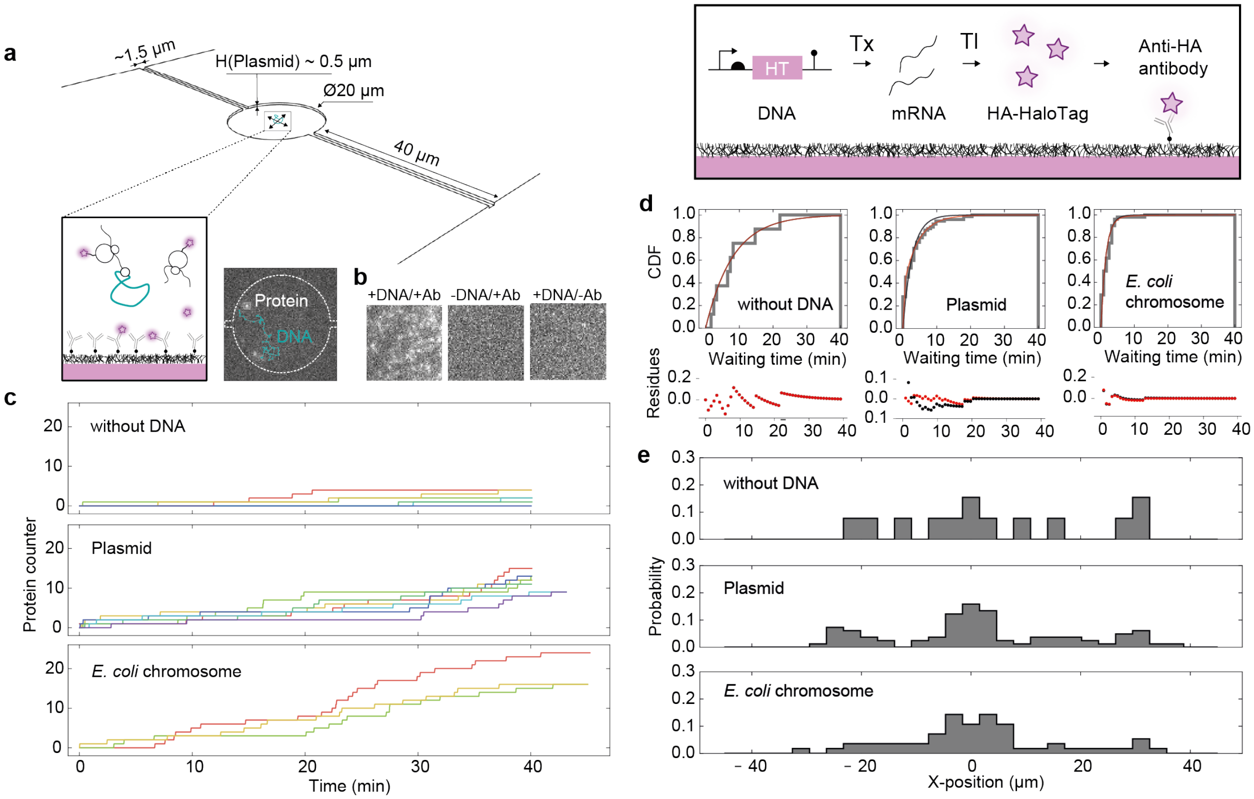


**Supplementary Figure 7: The production and capture of HTs from single DNA molecules.**

**a)** Schematic of a cell-like compartment with a freely diffusing plasmid encoding a cassette of native *E. coli* promoter, ribosomal-binding site, and HA-tagged HaloTag (HT) protein. RNAP transcribes the gene, and ribosomes translate the mRNA to produce HT proteins. The HA-HT proteins were captured on an anti-HA antibody-coated coverslip. A fluorescence image with a cell-like compartment (outlined with white dashed lines) containing diffusive (Atto647N-labeled plasmid) and static (surface-captured HT proteins) single-molecule spots. **b)** Fluorescence microscopy images with and without free-floating plasmid (HA-tagged HT) in cell lysate and 50 nM fluorogenic dye after 40 min of expression on the microfluidic chip outside the compartments. Only a few spots were detected when DNA or anti-HA antibodies (Ab) were omitted from the surface. **c)** Protein counter during cell-free expression in different cell-like compartments without DNA (n=6), with plasmid (n=7), and with proteinated chromosomes (n=3). **d)** The cumulative distribution functions of protein arrival times in cell-like compartments. The data was pooled from all experiments and fitted with a single (black curve) and double (red curve) exponential distribution to measure a potential deviation (residues are plotted in the lower panels) from random protein arrivals (=single exponential). **e)** Positional distribution of identified HT protein spots along the cell-like compartments. The capillaries were orientated along the X-axis.

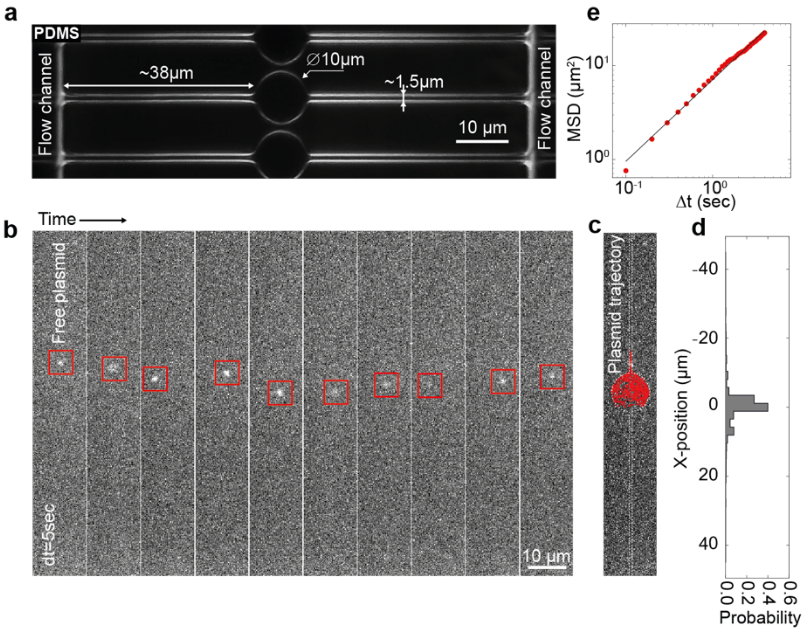


**Supplementary Figure 8: Embedding short DNA molecules in cell-like compartments.**
**a)** A darkfield image of cell-like compartments fabricated from polydimethylsiloxane (PDMS) to measure the diffusion coefficient of plasmids. Two capillaries connected cell-like compartments (diameter of 10 µm) with two main flow channels. The height of the compartment and capillaries were ~0.5 µm. **b)** Snapshots of a diffusing Atto647N-labeled plasmid in a cell-like compartment. **c)** The plasmid's trajectory (red line) was produced by single-molecule tracking. The dashed white line outlines the cell-like compartment. **d)** The probability distribution of plasmid positions in the compartment and capillaries. The compartment was centered at zero. **e)** The mean-square displacement (MSD) curve of a single plasmid (red points) and fit to normal diffusion (black line, D=1.7 $\mu m^2\ sec^{-1}$).

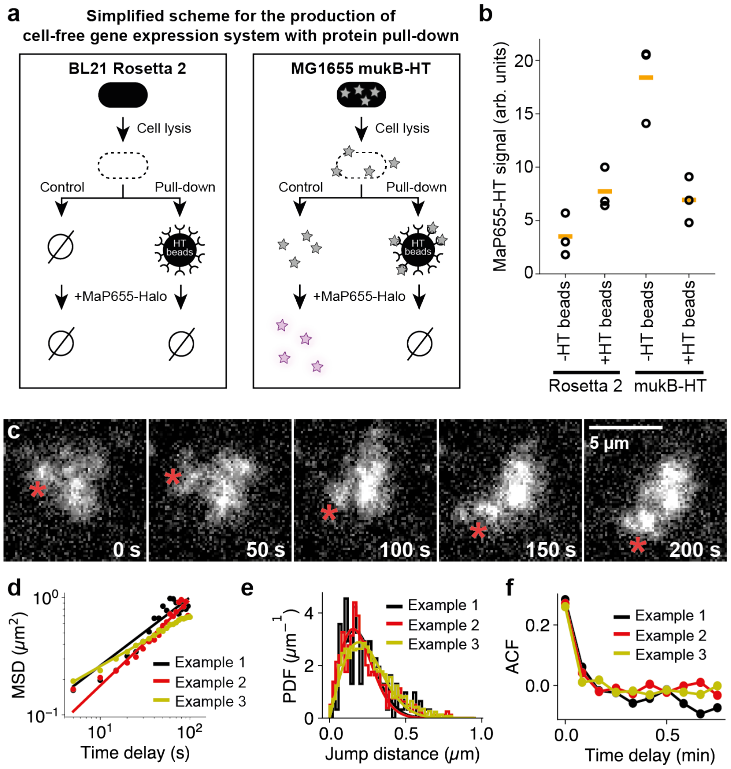


**Supplementary Figure 9: Spatial fluctuations on deproteinated chromosomes during cell-free gene expression.**

**a)** A schematic protocol for the pull-down of MukB proteins fused to HaloTag while preparing a cell-free gene expression system. We produced two cell lysates incubated with and without HaloTag immobilization beads (Magne HaloTag) during the run-off reaction. The cell-free gene expression system was incubated with 50 nM fluorogenic dye MaP655-Halo to estimate the mukB-HT levels in BL21 Rosetta 2 (neg. control) and MG1655 mukB-HT. **b)** The MaP655-Halo signal was measured with a well plate reader after incubation with the four lysate preparations. The yellow bars show mean values, and the black circles show data points from three technical replicates. **c)** An exemplary time-lapse of a single deproteinated chromosome diffusing in a large compartment during cell-free gene expression. We used persistent blobs (red asterisks) to track their locations and extract jump distances. The chromosome was labeled with SYBR Green I. **d)** The mean square displacement (MSD) curves for identified DNA blobs on three deproteinated chromosomes during cell-free expression. The three solid lines show the fits to a diffusion model $MSD = 4D\tau^n$ with $D$ as the apparent diffusion constant, $\tau$ as time delay, and $n$ as correction factor for a deviation from the normal diffusion model (n=1), producing apparent diffusion coefficients $D_1$=0.017±0.006 $\mu m^2\ s^{-1}$ (n=0.58), $D_2$=0.008±0.003 $\mu m^2\ s^{-1}$ (n=0.73), $D_3$=0.024±0.001 $\mu m^2\ s^{-1}$ (n=0.44). **e)** The probability distribution function (PDF) of jump distances $r$ of DNA blobs within five seconds on three deproteinated chromosomes during gene expression. The three solid lines show fits to a diffusion model (75) $P = N\, r/2D\tau\, e^{-r^2/(4D\tau)}$ with $N$ as a normalization factor, $\tau$ as the 5-second time delay between jumps, producing apparent diffusion coefficients $D$ of $D_1$=2.7e-3 ± 2.7e-4 $\mu m^2\ s^{-1}$, $D_2$=2.4e-3 ± 1.6e-4 $\mu m^2\ s^{-1}$, $D_3$=3.7e-3 ± 1.5e-4 $\mu m^2\ s^{-1}$. **f)** The median auto-correlation functions (ACF) of jump distances averaged over the identified tracks.

**Supplementary Table 1: Partial plasmid sequences**

| Name | Sequence (RBS is highlighted in red) |
|---|---|
| *N-terminal HA tag for HT (HA tag highlighted with bold letters until HT's atg)* | gcAATAATTTTGTTTAACTTTAAGAAGGAGATATAcc**ATGACCAGCTACCCATACGATGTTCCAGATTACGCTGGCCGCTTAATTAAACATATGACC**atg… |
| *Promoter (small letters in front of RBS) and RBS region up to start codon for the HUα-GFP plasmid* | gctgtgagcggataacattgacattgtgagcgggataacaagatactgagcacagctagcAATAATTTTGTTTAACTTTAAGAAGGAGATATAccatg… |

**Supplementary Table 2: Primers for *E. coli* genome engineering and plasmid construction**

| Name | Sequence (RBS is highlighted in red) | Comments |
|---|---|---|
| *MG1655-pKD4.f (capital letters = homolog region)* | GTCAGTTTAAATTATAAAAATTGCCTGATACGCTGCGCTTATCAGGCCTAgaaagcaggtagcttgcagtg | Forward primer to generate PCR fragment from pKD4-HA_HT for insertion into the chromosome |
| *MG1655-pKD4.r (capital letters = homolog region)* | TCCTGCGCTTTGTTCATGCCGGATGCGGCTAATGTAGATCGCTGAACTTGccatatgaatatcctccttagttcc | Reverse primer to generate PCR fragment from pKD4-HA_HT for insertion into the chromosome |
| *HA_HT.chr.f* | AAA TTG CCT GAT ACG CTG CG | Forward primer to verify chromosome integration of the HA_HT cassette and sequence |
| *HA_HT.chr.r* | GCA CCA GTA CGT TTT CCG CA | Reverse primer to verify chromosome integration of the HA_HT cassette and sequence |
| *hupA.f* | gcAATAATTTTGTTTAACTTTAAGAAGGAGATATAccatgaacaagactcaactgattgatg | Forward primer to lift the *hupA* gene from the *E. coli* genome and insert it into the pBEST backbone |
| *hupA.r* | AAGCTCCATGCTGGTCCCGGGAGCTCGCTTcttaactgcgtctttcagtgc | Reverse primer to lift the *hupA* gene from the *E. coli* genome and insert it into the pBEST backbone |
